# Molecular profiling of driver events and tumor-infiltrating lymphocytes in metastatic uveal melanoma

**DOI:** 10.1101/742023

**Authors:** Joakim Karlsson, Lisa M. Nilsson, Elin MV Forsberg, Suman Mitra, Samuel Alsén, Ganesh Shelke, Vasu R Sah, Ulrika Stierner, Charlotta All-Eriksson, Berglind Einarsdottir, Henrik Jespersen, Lars Ny, Per Lindnér, Erik Larsson, Roger Olofsson Bagge, Jonas A. Nilsson

## Abstract

Uveal melanoma (UM) is a rare form of melanoma with a genetics and immunology that is different from skin melanoma. Previous studies have identified genetic driver events of early stage disease when the tumor is confined to the eye. However due to lack of a clinical rationale to biopsy metastatic disease, access to tumor material to perform molecular profiling of metastases has been limited. In this study, we have characterized genomic events in UM metastases using whole-genome sequencing of fresh frozen biopsies from thirty-two patients and profiled the transcriptomes of individual tumor infiltrating lymphocytes in eight patients by single-cell sequencing. We find that 91% of the patients have metastases carrying inactivating events in the tumor suppressor *BAP1* and this coincided with somatic alterations in *GNAQ*, *GNA11*, *CYSLTR2*, *PLCB4*, *SF3B1* and/or *CDKN2A*. Mutational signature analysis revealed a rare subset of tumors with prominent signs of UV damage, associated with outlier mutational burden. We study copy number variations (CNV) and find overrepresented events, some of which were not altered in matched primary eye tumors. A focused siRNA screen identified functionally significant genes of some of the segments recurrently gained. We reintroduced a functional copy of *BAP1* into a patient-derived *BAP1* deficient tumor cell line and found broad transcriptomic changes of genes associated with subtype distinction and prognosis in primary UM. Lastly, our analysis of the immune microenvironments of metastases revealed a presence of tumor-reactive T cells. However, a majority expressed the immune checkpoint receptors TIM-3, LAG3 and TIGIT, and to a lesser extent PD-1. These results provide an updated view of genomic events represented in metastatic UM and immune interactions in advanced lesions.

## Introduction

Uveal melanoma (UM) is a rare form of melanoma but the most common intraocular cancer^1^. Enucleation or brachytherapy can provide good local control but in 50% of patients metastases develop, most frequently to the liver and generally with lethal outcome^1^. The genetics of UM has primarily been studied in the primary tumors of the eye, including that of the TCGA consortium^2^. Recurrent mutations in *GNAQ* or *GNA11* are common, whereas mutations in *PLCB4* and *CYSLTR2*, downstream and upstream of *GNAQ*/*11*, are seen in occasional cases^3–6^. These driver mutations are all mutually exclusive. Additional recurrent mutations have been found in *EIF1AX*, *SF3B1* and *BAP1*, where the latter connotes poor prognosis and development of metastatic disease^7, 8^. The development of metastatic UM can also be predicted using gene expression analyses, where Class 1 tumors have an excellent prognosis whereas Class 2 have a very poor prognosis, in a close to binary manner^9^.

Patients with UM metastases are not predicted to respond to the same targeted therapies as patients with cutaneous mutations since UM does not have *BRAF* mutations. Moreover, retrospective analyses of outcome following the use of immune checkpoint inhibitors have demonstrated poor response rates at multiple centers^10^. At our center, we are using isolated hepatic perfusion with melphalan to treat patients with liver metastases of UM. Retrospective analyses have suggested a survival benefit of this surgical method but this is now being challenged in a prospective randomized phase 3 trial (the SCANDIUM trial)^11^. Notably, during the surgical procedure leading to the perfusion treatment, there are possibilities of procuring fresh biopsies for the generation of PDX models, TIL cultures and for genomics studies of metastases (**Fig. 1a**). Here we describe a profiling of thirty-two metastatic UM tumors using whole-genome sequencing and we characterize infiltrating lymphocytes, providing molecular insight into the genomic events and immunogenetics driving late-stage UM.

**Figure 1.**
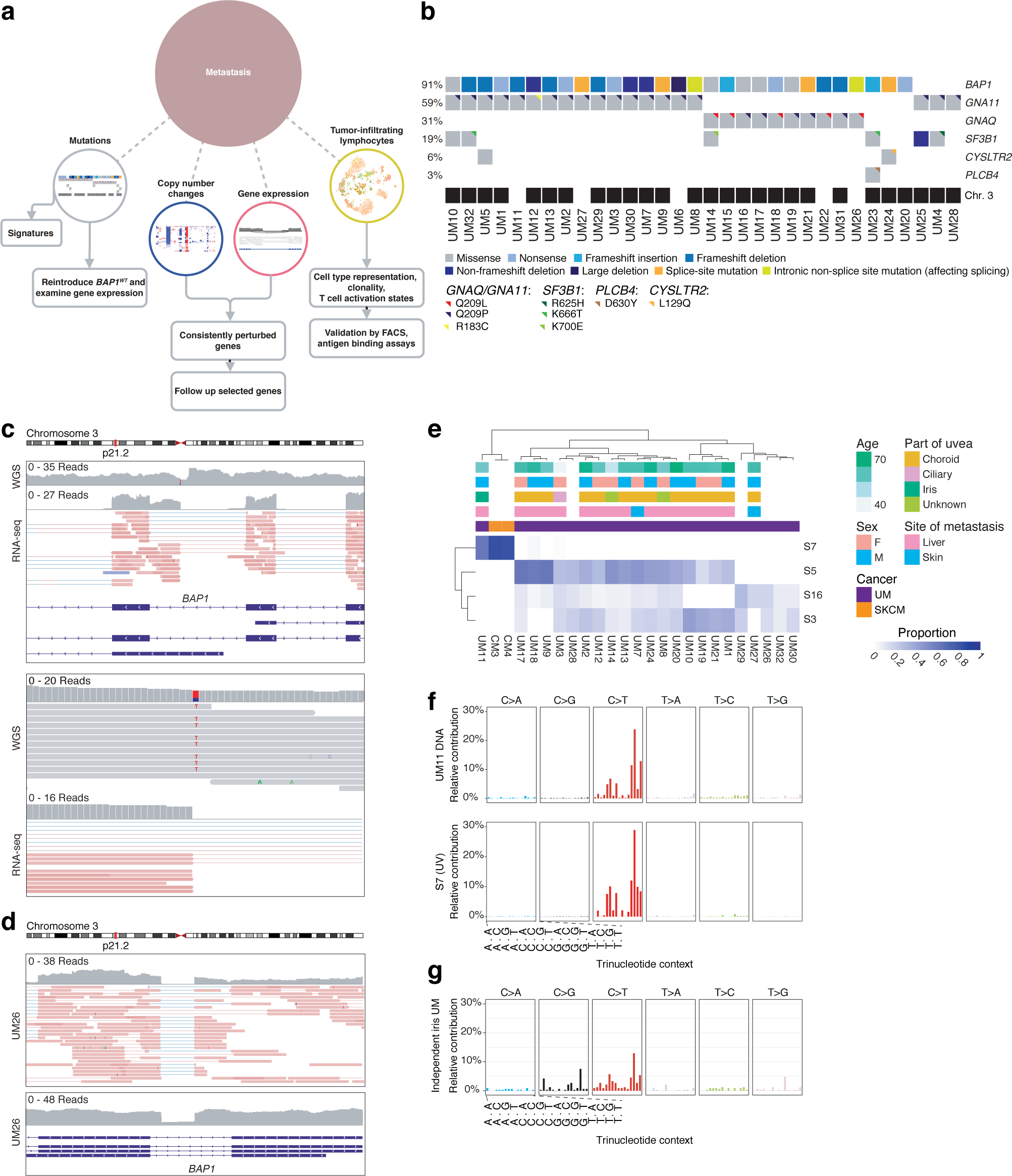
Recurrently mutated genes and copy number aberrations in metastatic uveal melanoma (UM). **a)** Schematics of the study. **b)** Mutations in genes recurrently altered in UM. Chromosome 3 status is indicated. **c)** Intronic non-splice site point mutation in *BAP1*, associated with aberrant splicing. **d)** Intronic large deletion in *BAP1* associated with aberrant splicing. **e)** Estimated contributions of COSMIC mutational signatures. Samples and signatures are ordered by agglomerative hierarchical clustering. Signatures with estimated contribution < 25% excluded. Two cutaneous melanomas (indicated “CM”) sequenced at the same time were included for comparison. Only tumors with matching normals were included. Signatures were inferred using both synonymous and non-synonymous mutations in exonic regions. **f)** Overall mutational spectrum of UM11, shown on WGS and RNA-seq data. The canonical profile of the UV-associated “signature 7” is shown for comparison. **g)** Mutational spectrum of an unrelated iris melanoma of a primary site. The tumor was sequenced by exome sequencing.

## Results

### Recurrently mutated genes in UM metastases

Thirty-two metastases of UM, six subcutaneous and 26 from the liver (**Table S1**), were collected and subjected to whole-genome sequencing (WGS) and 26 of them to poly-A+ RNA sequencing. Twenty-eight of the tumors were pathologically designated as originating from the choroid, one in the ciliary body and one in the iris, while two cases were ambiguous. All liver metastases came from patients that were untreated at the time of biopsy and 24 of them had been enrolled in the SCANDIUM phase III trial. All cutaneous biopsies except one came from patients previously treated with chemotherapy (IHP, dacarbazine and/or taxanes).

Variant calling with MuTect 2^12^ revealed mutations in *BAP1*, *GNA11*, *GNAQ*, *SF3B1*, *CYSLTR2*, and *PLCB4* (**Fig. 1b**, **S1** and **Table S2**), which are recurrently altered in UM^2–4, 7^. We discovered no mutations in *EIF1AX*, which is associated with a good prognosis^2, 7^. In all, 29/32 (91%) of metastases were found to harbor *BAP1* mutations. These were paired with loss of chromosome 3 the vast majority of cases (**Fig. 1b**). Notably, *BAP1* was also the subject of alterations not detected by standard variant calling, including one large deletion spanning the first three exons. In another case, one intronic event far from the nearest splice site was associated with novel splicing events at the point of the mutation and intron retention (**Fig. 1c**). A third tumor contained a 48 bp fully intronic homozygous deletion that again did not occur at a splice site, but associated with mis-splicing and intron retention clearly tied to the event (**Fig. 1d**). These two alterations most likely created new intronic splice sites. A previous study has described a mutation that activates a cryptic splice site within an exon in *BAP1*^13^. To our knowledge no cases have been described for de novo splice-site-generating intronic mutations in UM; only cases that disrupt canonical splice sites at the exon-intron boundary^14^. Since *BAP1* loss predicts metastasis^15^, this highlights the need to also investigate intronic non-splice site mutations as candidates for loss-of-function events, which exome^2^ or targeted^16^ sequencing may not be sufficient to reveal.

Among the three patients where *BAP1* mutations could not be established, two had *SF3B1* mutations. We also detected mutations in *SF3B1* that occurred outside the common hotspots K666 or R625. These were K700E, I955S and an in-frame deletion at V577. The first has to our knowledge not been described in UM, but is frequent in other cancer types, including breast cancer^17^, chronic lymphocytic leukemia^18^ and pancreatic adenocarcinoma^19^. Some *SF3B1* mutations also co-occurred in tumors with *BAP1* mutations, illustrating that mutual exclusivity between these events is not perfect^2, 3^.

In the third tumor without *BAP1* mutation, we did not discover mutations in either *SF3B1* or *EIF1AX*. This tumor (UM28) was also the only one inferred to be tetraploid (**Fig. S2b**), and had frequent wide copy number losses, affecting chromosomes 1p, 3, 4q, 6q, 8p, 9, 11, 14 and 16. Mutated genes in these regions included *YEATS2* and *ZMAT3* on chromosome 3 and *AKT1* on chromosome 14. This sample also displayed the second highest levels of *PRAME* expression, which has been independently associated with metastasis^20^.

In addition, we found two metastases with mutations in the tumor suppressor *TET2*, in one case leading to a stop-gain. A third tumor had a frame-shift deletion in *TET1*. *TET1* and *TET2* exert epigenetic control via DNA demethylation^21, 22^. Some metastases also had mutations in genes that interact with *BAP1*, including *ASXL2* and *FOXK2*^23^ (**Table S2**).

### Mutational signature of UV damage in UM

The causes that underlie UM are to date largely unknown, and despite risk factors implying a potential role for UV radiation, no clear evidence has emerged to date and the field is divided on whether this can be a driving factor^1, 4, 24–28^. The pattern of trinucleotide substitutions across the genome can be informative about underlying mutational processes. Therefore, we estimated the relative contributions of established mutational signatures^29^ to the total exonic mutational burden in the tumors.

Consistent with previous observations^4^, the dominating signatures were S3, S5 (COSMIC nomenclature), and to a lesser extent S16. S3 has been associated with defective DNA double-stranded break repair, whereas S5 is termed “clock-like” and associates with aging^29, 30^ (**Fig. 1e**). However, one tumor had a distinctly different profile, dominated by contributions from S7 (about 63%), with a bias towards the untranscribed strand (*q* < 0.05, Poisson test), more closely resembling cutaneous melanomas sequenced concurrently (**Fig. 1e**-**f**). S7 is known to arise as a consequence of UV radiation-induced damage^29^. We could exclude a mix-up from the presence of the same *GNA11* Q209L mutation and *BAP1* frame-shift deletion in RNA, together with transcriptomic classification against ∼10000 tumors from TCGA (**Fig. S3a-c**).

We hypothesized that this unexpected signature could be explained due to the tumor having originated in the iris (**Fig. S3d-e**), a site from which only 3-5% of cases arise^31^, compatible with an absence of iris melanomas and UV evidence in the TCGA UM cohort^2^. To confirm this, we managed to obtain a second iris UM sample from a patient without metastasis, which again revealed a prominent a UV pattern (**Fig. 1g**). Thus, although rare, UM can evidently be induced by UV damage if manifest in the iris.

### Copy number changes overrepresented among metastases

UM is characterized by highly recurrent copy number aberrations affecting entire chromosome arms^2^. All metastases had gain of chromosome 8q, known to co-occur with monosomy 3 in poor prognosis tumors^26, 32, 33^ (**Fig. 2a**). A number of arm-level changes were also significantly overrepresented in the metastatic tumors compared to tumors studied by TCGA (Fisher’s exact test, *q* < 0.05). These were loss of 17p and 6q, as well as gain of 8q and 5p gain (**Fig. 2a**-**b**, **Table S3**). Loss of chromosome 3 was close to significance at *q* < 0.094. Previous studies have also found loss of 6q and 8p to be overrepresented in metastatic tumors^26, 32^. Sequencing of matched primary tumors for UM16 and UM24 showed that 8q gain and loss of 3 was present already in the primaries in both cases (**Fig. 2c**). However, gain of 5p in UM16 and loss of 6q in UM24 were later events only present in the respective metastases. Overall, genomic losses tended to be more frequent in these metastases than observed in TCGA tumors.

**Figure 2.**
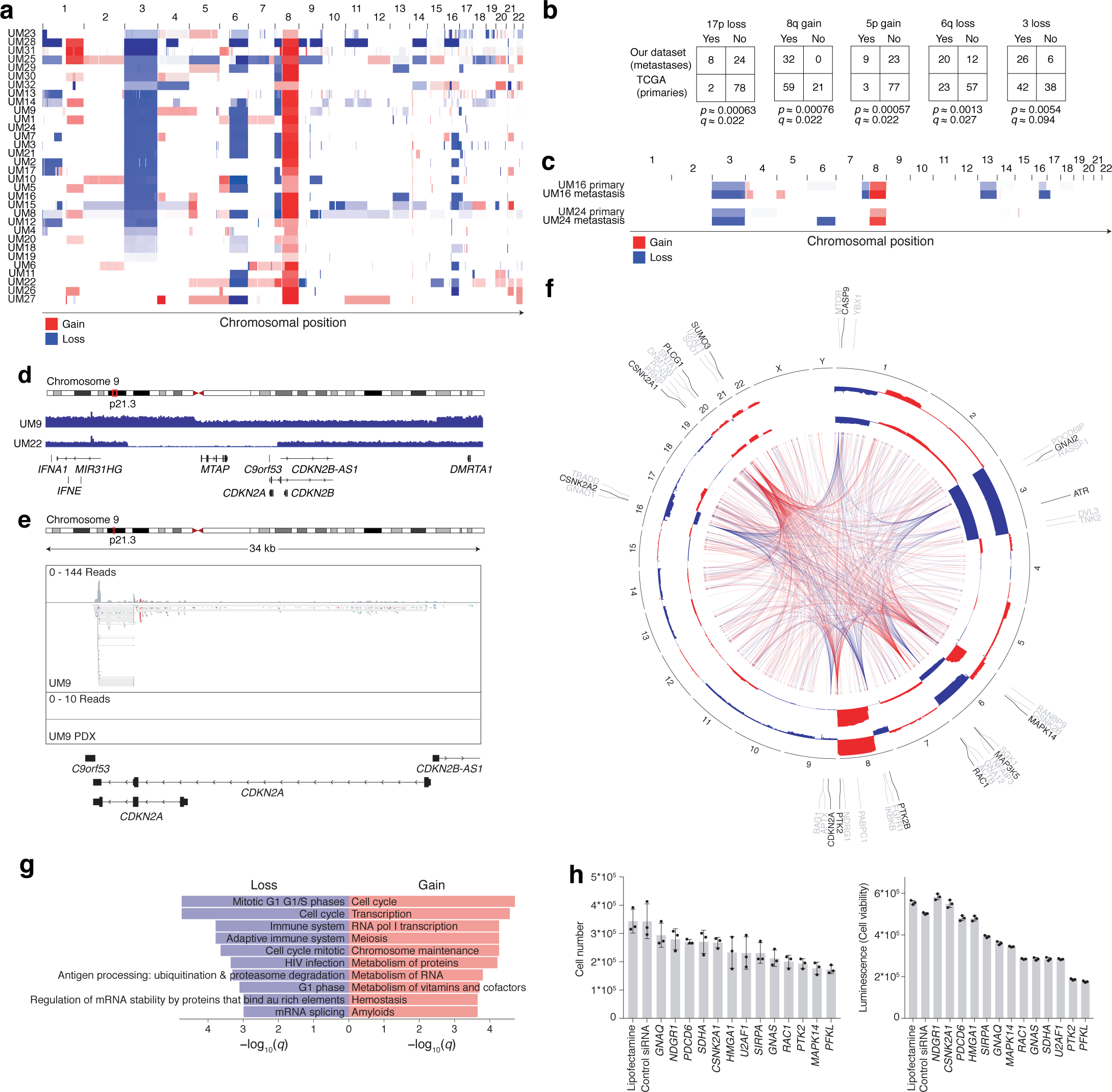
Copy number analysis. **a)** Copy number profiles of each tumor. **b)** Broad copy number changes enriched in the metastases (*n* = 32) compared to TCGA tumors (*n* = 80), assessed using two-tailed Fisher’s exact tests and adjusted for multiple testing using the Benjamini-Hochberg method. **c)** Copy number profiles of two primary tumors together with the corresponding matched metastases. **d)** Focal deletions of *CDKN2A* in two samples. **e)** RNA-seq of the samples with focal *CDKN2A* deletions in samples taken from the metastases and samples taken from PDX models established from these tumors. **f)** Genes in recurrent arm-level copy number aberrations ranked by correlation between gene expression and copy number consistent among the metastases and in TCGA tumors as well as protein-protein interaction network degree from the Human Protein Reference Database (HPRD), with the top three candidates shown in each region. Connecting lines represent protein interactions of the highest ranked gene per region. Blue represents regions of loss and red regions of gain. Summarized representations of copy number profiles per region show the relative numbers of gain and loss events, with the inner circle representing TCGA samples and the outer our cohort. **g)** Gene pathways enriched among the combined set of genes per region of gain or loss. **h)** Functional interrogation by siRNA of a selected amount of genes whose expression is elevated due to CNV. Cells were counted or viability were measured 72 h after transfection of the siRNA pools.

Focal events were very rare. Notably, however, we discovered somatic focal deletions affecting *CDKN2A* and the nearby gene *MTAP* in two samples (**Fig. 2d**, **Fig. S4**). *CDKN2A* encodes the tumor suppressors p16^INK4a^ and p14^ARF^ and is commonly deleted in cutaneous melanoma^34^. The deletions here were homozygous and hemizygous respectively. While *CDKN2A* expression was still present in the hemizygous case, a subsequent patient-derived xenograft (PDX) model established from this tumor showed full loss of expression, even extending to other nearby genes (**Fig. 2e**, **Fig. S5**). This suggests that either a pre-existing clone with a homozygous deletion or a second loss event was selected for as the tumor established itself in this new environment, supporting *CDKN2A* loss as a late event that may be relevant in the metastatic setting.

### Survey of genes in recurrent arm-level copy number events that may influence tumor behavior

To understand how the recurrent chromosomal events in UM affect the transcriptome and to rank genes by a potential to influence tumor behavior, we searched for consistent correlations between the copy number of each gene affected and its expression in this dataset and TCGA UM, and ordered them by their degree of known protein-protein interactions from the HPRD database, followed by association with survival. The top candidates per region are shown in (**Fig. 2f**, **Table S4**). An analysis using the “chemical and genetic perturbations” collection in MSigDB showed that regions of gain were enriched for the category “uveal melanoma class II up” (*q* < 1.84*10^-19^), whereas regions of loss were enriched for “uveal melanoma class II down”^35^ (*q* < 3.15*10^-17^). The class II transcriptional subtype is one of the two major subdivisions of UM, strongly associated with metastasis^35^. A Reactome enrichment analysis revealed processes that included cell cycle progression, chromosome maintenance, immune signaling and hemostasis (**Fig. 2g**, **Table S5**).

Top ranked genes in loss regions included *CASP9*, an early activator of apoptosis^36^ and the aforementioned *CDKN2A*. Candidates in gain regions included *MAPK14* (p38α), a kinase that operates at the intersection of cell cycle progression, stress signaling, immune responses and differentiation^37–40^, and the very recently proposed UM oncogene *PTK2* (FAK)^41^, a negative regulator of cell detachment-initiated apoptosis (anoikis)^42, 43^. A small RNAi screen, directed against a list of genes selected based on gain candidates, in a cell line derived from the UM22 tumor demonstrated that 8/12 siRNA pools negatively affected proliferation (cell count) or viability (ATP production) to a similar or higher level than an siRNA against *GNAQ* (**Fig. 2h**). Thus, these recurrent arm-level copy number changes contribute to shaping the transcriptomic subtypes of UM and regulate genes that may conceivably contribute a fitness advantage.

### *BAP1* loss contributes to a transcriptomic shift towards the metastatic class II subtype and up-regulates TIM-3 and TIGIT immune checkpoint ligands

We next asked to what extent *BAP1* loss could influence the transcriptome of metastatic UM. For this purpose, we used the UM22 cell line, which had been established from one of the metastases grown as a PDX, which had a homozygous frame-shift deletion in *BAP1* (**Fig. S5**). A functional copy of *BAP1* was introduced using a retroviral vector and RNA-seq performed on this and an empty vector control sample (**Fig. 3a**). RNA-seq alignments showed successful integration of the wild-type *BAP1* allele (**Fig. 3b**). A differential expression analysis between the two conditions revealed a large transcriptomic response, with 518 genes downregulated and 990 upregulated at an absolute log_2_ fold change > 1 and *q* < 0.05 (**Fig. 3c**, **Table S6**). *SLC7A11*, identified by Zhang et al. as a mediator of ferroptosis-suppressive effects of *BAP1*^44^, was significant albeit not as strongly regulated (log_2_ fold change = −0.82, *q* = 5.42*10^-19^). Pathways enriched among downregulated genes upon reintroduction included GPCR signaling, neurotransmitter receptor transmission, interferon alpha/beta signaling and chemokine activity. Upregulated pathways most prominently included post-transcriptional and translational mechanisms (**Fig. 3d**).

**Figure 3.**
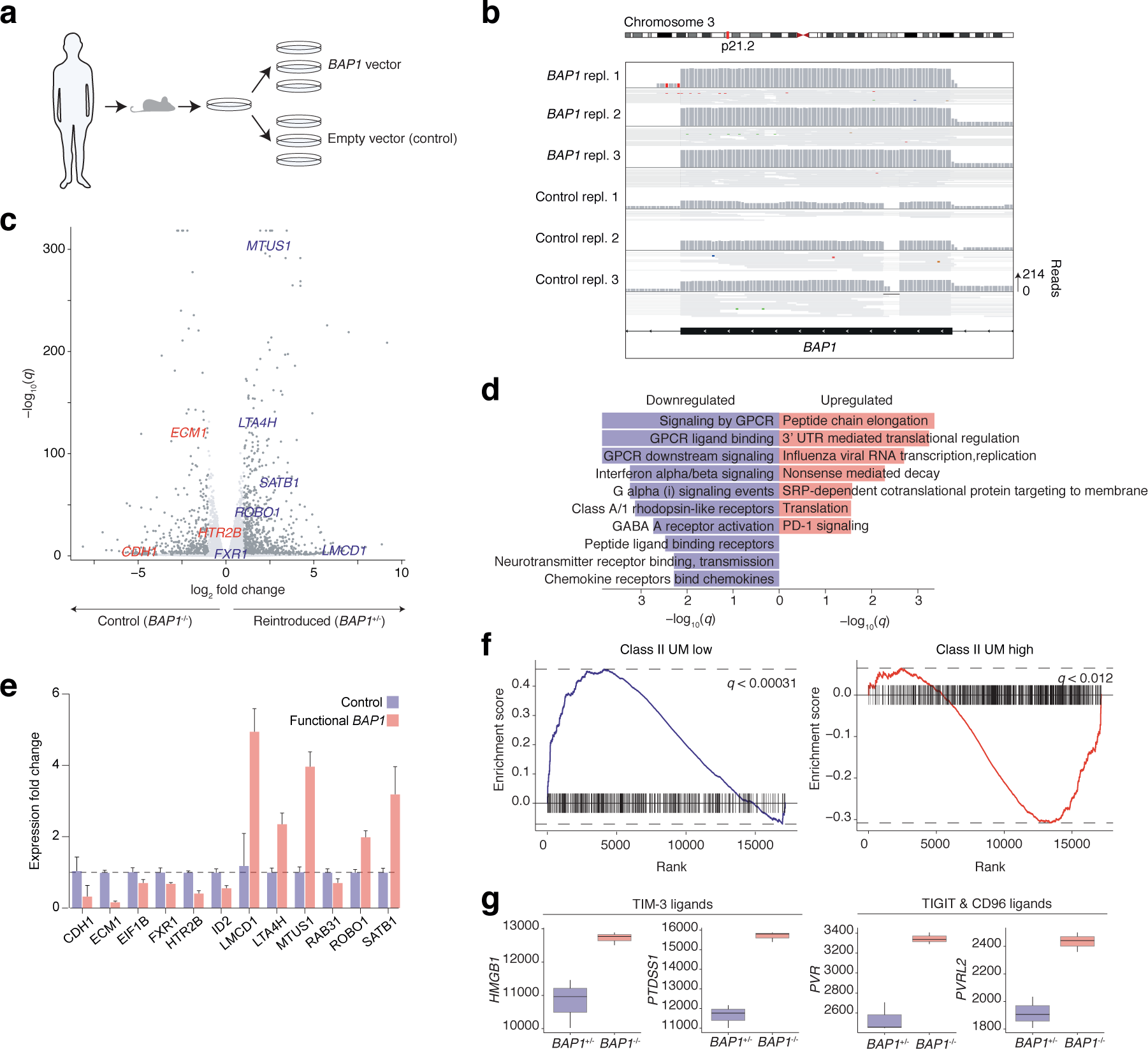
Reintroduction of *BAP1* into a deficient tumor. **a)** Schematic representation of the experiment. Cell lines from a PDX model established from tumor UM22 were transduced with either *BAP1* wild-type containing viral vectors or empty vectors and subjected to RNA sequencing. **b)** RNA-seq alignments of *BAP1* in the two conditions. **c)** Differentially expressed genes for *q* < 0.05 and absolute log_2_ fold change > 1. *n* = 3 biological replicates were used and differences were assessed using DESeq2, with the parameter alpha = 0.05. Genes from a clinical assay distinguishing the class I and II UM subtypes are indicated^45^. **d)** Top ten enriched gene sets for the “canonical pathways” MSigDB category. **e)** RT-qPCR results for the genes indicated in (c), with *n* = 3 biological replicates. Bars represent means and error bars represent standard deviations. **f)** Gene set enrichment analysis with respect to MSigDB “chemical an genetic perturbation” category, with results from the two sets discriminating between class I and II subtypes shown^35, 47^. **g)** Downregulation of ligands to the immune checkpoint receptors *TIM-3*, *TIGIT* and *CD96* upon *BAP1* reintroduction.

Notably, we observed significant regulation of nine out of 12 genes used as discriminating features in a classifier that distinguishes between the high-risk class II versus class I subtypes^45, 46^, some of which are melanocyte lineage markers and a few of which have also been found compatibly regulated upon silencing^8^ (**Fig. 3c**). These genes were all expressed in the inverse fashion expected for class II tumors, with *CDH1*, *ECM1* and *HTR2B* decreasing upon *BAP1* reintroduction and *LMCD1*, *LTA4H*, *MTUS1*, *ROBO1*, *SATB1* and *FXR1* increasing. This was confirmed with additional RT-qPCR measurements for all genes but *FXR1* (**Fig. 3e**).

To investigate whether this trend was limited to these few discriminating genes or representative of a broader transcriptomic shift towards the class I subtype, we performed a gene set enrichment analysis on the whole list of differentially expressed genes using the “chemical and genetic perturbations” collection from MSigDB^47^. We found the “uveal melanoma class II down” gene signature^35^ to be significantly enriched among upregulated genes (*q* < 0.00031), and “uveal melanoma class II up” to be enriched among downregulated genes (*q* < 0.012), showing that this trend is indeed broader (**Fig. 3f**, **Table S7**). This shift towards the class I subtype upon *BAP1* reintroduction implies that the inverse drives the cells towards the metastatic class II transcriptional subtype, which characteristically has *BAP1* alterations.

Beyond this, *BAP1* restoration also downregulated the TIM-3 immune checkpoint ligands *HMGB1* and *PTDSS1* as well as the *TIGIT* and *CD96* ligands *PVR* and *PVRL2*, implying higher expression levels in *BAP1*-deficient UM cells (**Fig. 3g**, **Table S6**).

### T cells from UM metastases recognize tumor antigens and predominantly express the checkpoint receptors LAG3, TIM-3 and TIGIT

Having observed the regulation of checkpoint proteins in UM cells, we next investigated the phenotypes of tumor-infiltrating lymphocytes (TILs) isolated from metastases. Despite the generally poor immunogenicity of UM, we could confirm the presence of melanoma-specific TILs in a subset of patients (**Fig. 4a**). This included the UV-associated iris tumor, which was also predicted to have the highest neoantigen load (**Fig. 4a** and **Fig. S6**). More detailed flow cytometry of cryopreserved single cell preparations of tumors revealed PD-1^+^CD39^+^ cells present in high fractions of CD8+ T cells in a subset of samples (**Fig. 4b-c**, gating strategy in **Fig. S7a-b**). PD-1^+^CD39^+^ cells have been proposed to be tumor-reactive^48^. Most samples maintained their relative proportions of CD8+ and CD4+ T cells after expansion (**Fig. S7c**). We therefore performed single-cell RNA and T cell receptor (TCR) sequencing of TILs from these eight tumors for a more comprehensive view.

**Figure 4.**
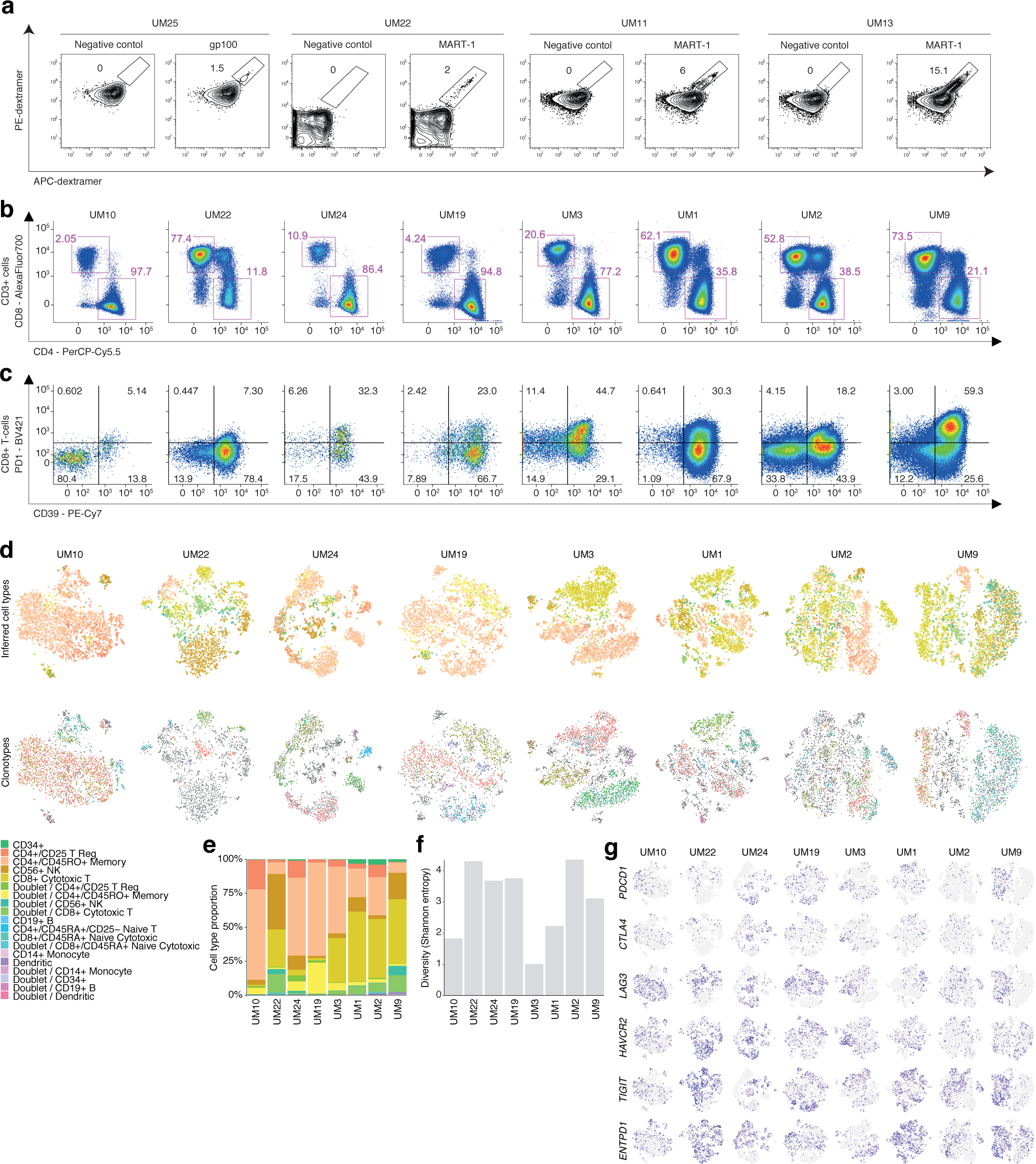
Analysis of tumor-infiltrating lymphocytes. **a)** Assessment of T cell reactivity against MART-1 and gp100. Proportions found to be specific are indicated. **b-c)** Flow cytometry analysis of T-cells. (a) Proportions of CD8+ and CD4+ cells. (b) Proportions of PD-1+ and CD39+ CD8+ cells. **(d-g)** Paired transcriptome and TCR profiling of T cells. **d)** t-SNE representations of cell transcriptomes, colored by inferred cell type (upper) and the most abundant T cell receptor clonotypes (lower). **e)** Relative proportions of each cell type. **f)** TCR diversity, measured by Shannon entropy. **g)** Expression of exhaustion markers in each cell.

Inference of cellular subtypes by correlation to pure PBMC subsets^49^ revealed similar proportions of CD8+ and CD4+ subsets as observed by flow cytometry (**Fig. 4b-d** and **Fig. S7d**). The transcriptome analysis also revealed heterogeneity in regulatory CD4+ T cell fractions, potentially suggesting samples with more suppressive mircoenvironments. Clusters were formed both by cell type and receptor clonotype (**Fig. 4d-e**), revealing clones in different activation states and clonal expansion (**Fig. 4f** and **Fig. S8**). Notably, we observed abundant expression of activation and exhaustion markers (**Fig. 4g** and for confirmation by flow cytometry, **Fig. S9-10**). The most prominently expressed checkpoint receptors were LAG-3, TIM-3 and TIGIT, with fewer cells expressing PD-1 and CTLA-4. The predominance of LAG-3, TIM-3 and TIGIT, together with the observation that *BAP1* can suppress ligands for the former two, indicates means of immune evasion in UM that are different from cutaneous melanoma.

## Discussion

Metastatic UM currently entails a very poor outcome due to the lack of effective treatment options^15^. Genetics of the primary disease confined to the eye has already been investigated in several hallmark studies^2–4, 7–9^. However, only a few metastatic samples have been sequenced with exome or whole-genome sequencing and our study has the largest sample cohort sequenced to date with whole-genome sequencing. A history of primary uveal melanoma and lack of therapeutic efficacy of surgery makes biopsy and surgical removal of samples not clinically meaningful. By obtaining biopsies of liver metastases from patients in the SCANDIUM trial or cutaneous metastases of UM, we have been uniquely positioned to focus on the metastatic disease by both analyzing fresh frozen material by genomics as well as generating PDX, cell lines and TIL cultures for transciptomics analyses.

The key event to metastasis in UM is loss of the tumor suppressor *BAP1*^8^. Compatible with this, we observed *BAP1* alterations in 91% of the metastases. Two of these altered splicing via intronic events outside of canonical splice regions, via creation of new intronic splice sites. This illustrates special cases that exome sequencing may not be sufficient for detecting, and which have high prognostic value.

We furthermore find that out of two tumors studied of the iris subtype, both had mutational spectra associated with UV-induced damage. Mutational signatures of UV damage in UM have not previously been reported and a consensus of UV-involvement in UM has not been reached by previous epidemiological studies. While iris UM is rare, the metastasis studied here had much higher than average mutation load, and predicted number of neoantigens. This could potentially render such tumors suitable for immunotherapy, which otherwise lacks efficacy in UM. Interestingly, the iris UM metastasis concerned here also harbored T cells recognizing MART-1.

Several broad copy number events were found to be more frequent in the metastases studied compared to primary tumors from TCGA, including losses of 17p loss and 6q, as well as gains of 8q and 5p. Notably, 8q gain was present in every metastasis. By sequencing matched primary tumors for two cases, we could establish that in one of the tumors 5p gain and 17p loss had arisen during metastasis, and in the other case 6q loss. Furthermore, two tumors had focal deletions of *CDKN2A*, an event that may have a larger relevance in the metastatic setting, as it has not been detected in recent large-scale studies of primary UM tumors^2–4, 26, 28^.

We additionally mapped out genes with correlations between expression and arm-level copy number changes in both this dataset and that of TCGA and ranked them by their degree of protein-protein interactions and any associations with survival present to gain an understanding for central processes affected and potential targets. We found several interesting candidates, including the recently proposed UM oncogene *PTK2*^41^, *MAPK14*, the apoptosis mediator *CASP9*^36^, as well as *CDKN2A* to be first-ranked candidates in 8q gain, 6p gain, 1p loss and 9p loss, respectively. We performed an siRNA knockdown experiment against selected genes and found proliferation and viability decreases to be the consequence when targeting the majority of those. In addition, we found expression changes mediated by loss events to be enriched for genes generally downregulated in poor-prognosis tumors and gain events enriched for genes upregulated in poor-prognosis tumors, showing that these broad events contribute to shaping the distinct transcriptomes of the two subtypes.

To increase our understanding for how these transcriptomic subtypes are established, we investigated the contribution from *BAP1* loss by reintroducing a functional allele into a cell line established from one of these metastases. We found that genes upregulated in cases with the functional gene where enriched for those that are lowly expressed in the poor-prognosis class II subtype, and vice versa. In essence, reintroduction indicated a reversal of the transcriptomic subtype. Notably, also immune checkpoint ligands where found regulated by *BAP1*. Of potential importance was the downregulation of TIM-3 and *TIGIT* ligands in the cases with functional *BAP1*, indicating a potential upregulation on *BAP1* loss that may have consequences for tumor-immune interactions.

We profiled the transcriptomes of tumor-infiltrating lymphocytes isolated from eight of the metastases by flow cytometry and single-cell sequencing and found tumor-reactive subsets present in several cases. However, their transcriptomes also indicated a high degree of exhaustion, with prominent expression of the checkpoint receptors LAG3, TIM-3 and TIGIT, and to a lesser extent PD-1 and CTLA-4. Potentially, the inferred upregulation of ligands for TIM-3 and TIGIT upon *BAP*1 loss may cooperate with the high level of expression of these receptors by T cells to interfere with anti-tumor immunity. Given the historic failures of anti-PD-1 and anti-CTLA-4 therapies in UM, this may argue for exploring these other checkpoint mechanisms.

Collectively, these results highlight that exome sequencing may not be sufficient to detect *BAP1* loss, the most significant event in UM metastasis, that UV damage underlies an important mutational process in the iris subtype and that recurrent copy number aberrations cooperate with *BAP1* loss to shape the transcriptome of the metastatic subtype. We also describe immune-profiles of T cells present in metastases that indicate tumor recognition, but exhaustion with predominant activation of checkpoints that are not targeted by current immunotherapies.

## Methods

### Processing of tumor biopsies

The patients received oral and written information and signed the informed consent according to the ethical approval at the Regional ethical review board (#289-12 and # 144-13). Biopsies were either extracted from subcutaneous metastases or from liver metastases, during the procedure of isolated hepatic perfusion in the SCANDIUM trial for those participating in it. Tumor biopsies were divided into pieces that were snap-frozen or minced and used for cryopreservation or tumor-infiltrating lymphocyte cultures. Primary eye tumors were formalin fixed and paraffin embedded (FFPE) in blocks at St Erik’s Eye Hospital’s pathology biobank.

### Sequencing

DNA and RNA from fresh frozen biopsies, blood and tumor-infiltrating lymphocytes were extracted using the AllPrep DNA/RNA kit (Qiagen). Primary eye tumors were sectioned and processed using an FFPE DNA kit (Qiagen). Libraries were made using Illumina TruSeq kit and sequenced on HiSeq2500 instruments at SciLifeLabs in Stockholm or on a NovaSeq at GeneCore SU in Gothenburg. Exome sequencing libraries were prepared with the Nextseq500 Kit HighOutput v2 and sequenced with Nextseq500.

### Preprocessing of DNA sequencing data

Raw whole-genome sequencing reads were aligned to the 1000 Genomes version of the GRCh37 reference genome with bwa^50^ (version 0.7.12; options “mem” and “-M”). Duplicates were marked with Picard (version 1.109; https://broadinstitute.github.io/picard). The resulting BAM files were recalibrated with GATK BaseRecalibrator (version 3.3.0)^51^, supplying lists of known polymorphic sites from dbSNP v138 and 1000 Genomes. PDX samples were aligned separately to the human reference genome and to the GRCm38 version of the mouse reference genome. Reads originating from human were then determined using Disambiguate (version 2018.05.03)^52^, specifying the parameter “-a bwa”.

### Preprocessing of RNA sequencing data

RNA sequencing reads were aligned to the 1000 Genomes version of the GRCh37 reference genome with STAR^53^ (version 2.7.1a) with the parameters “--twopassMode Basic -- outFilterType BySJout, --outSAMmapqUnique 60”. Splice junctions were provided from the Ensembl GRCh37.75 reference annotation. Gene expression was quantified using htseq-count^54^ (version 0.6.0), with parameters “-m intersection-strict -s reverse”. Transcript-level expression was quantified using kallisto^55^ (default parameters), based on cDNA sequences corresponding to the Ensembl annotation of the GRCh37 human reference genome. PDX samples were aligned separately to the human reference genome and to the GRCm38 version of the mouse reference genome with STAR. Reads originating from human were then determined using Disambiguate, specifying the parameter “-a star”.

### Variant calling

Variant calling was performed with MuTect 2 (GATK v. 4.0.11.0) with a panel of normals for paired tumor and normal samples and in a minority of cases on tumor samples alone, with lists of known population variants provided from the Genome Aggregation Database^56^, specifying the parameters “--af-of-alleles-not-in-resource 0.0000025” and “--disable-read-filter MateOnSameContigOrNoMappedMateReadFilter”. Construction of a panel of normals was done by first running MuTect2 in on each normal with the parameter parameter “-- disable-read-filter MateOnSameContigOrNoMappedMateReadFilter” and then merging the resulting lists with “CreateSomaticPanelOfNormals” from GATK. MuTect2 calls were filtered by first running “FilterMutectCalls” and then removing all that failed these filters. Variant annotation was performed with VEP (v. 91.3) and ANNOVAR^57^ (version 2016-05-11), using the databases COSMIC (v. 79), ESP6500 (v. “siv2_all”), 1000 Genomes (v. “2015aug_all“) and dbSNP (v. “snp138NonFlagged”). For two of the tumors, exome-sequenced normals were used for further filtering using GATK SelectVariants.

### Mutational signature analysis

To determine mutation spectra, all exonic somatic mutations (including synonymous) not present in any population variant resource were converted into a 96-trinucleotide mutation frequency matrix using the function “mut_matrix” (parameter: ref_genome = “Bsgenome.Hsapiens.UCSC.hg19”, excluding the sex chromosomes) from the R package MutationalPatterns^58^. Known mutational signature trinucleotide frequencies, obtained via COSMIC (http://cancer.sanger.ac.uk/cancergenome/assets/signatures_probabilities.txt; accessed October 27, 2017), were then fitted to the observed mutations using the function “fit_to_signatures” of the same R package. This algorithm operates by searching for the nonnegative linear combination of the predefined mutational signatures that best explains all mutations in a given sample, which is done by solving a nonnegative least squares optimization problem^58^. As a result, estimations of the relative contributions of the known mutational signatures in each sample were obtained.

### HLA-genotyping

HLA genotyping was performed using polysolver (version 1.0)^59^ on whole-genome sequencing data, with the parameters “Unknown 0 hg19 STDFQ”.

### Neoantigen prediction

Mutated 17-mer peptide sequences centered at each mutation were constructed from non-synonymous point mutations not present in any population variant resource. Neoantigen predictions against the HLA class I genotypes of each sample were then performed using netMHCpan^60^ (version 4.0, default parameters), considering only 9-mers. Peptides with predicted affinity < 500nM were retained and those deriving from transcripts without expression were removed.

### Generation of PDX models and a *BAP1* deficient UM cell line

Animal experiments were performed in accordance with E.U. directive 2010/63 (Regional animal ethics committee of Gothenburg approval #36-2014). Cryopreserved biopises were thawed and single cells were transplanted into the flank (UM22) or the liver (UM9) of immunocompromised, non-obese severe combined immune deficient interleukin-2 chain receptor γ knockout mice (NOG mice; Taconic, Denmark) to form xenografts. Tumors were analyzed by immunohistochemistry using clinically used antibodies against Melan-A, PMEL (HMB45) and S100. For generation of a cell line, a PDX tumor was minced and seeded at high density into a 5 cm culture plate in RPMI medium supplemented with 10% fetal bovine serum. Surviving cells were expanded and characterized by RNA-seq. The cells were transduced with a retrovirus expressing HA-tagged BAP1 or a control retrovirus (MSCV-IRES-GFP), both of which were made using plasmids from Addgene.

### Differential expression analysis

RNA-seq data was aligned and quantified as described. Differential expression was assessed using DESeq2, with the parameter “alpha=0.05”. Genes with *q*-values below 0.05 were considered statistically significant. Gene set enrichment analysis was carried out with the R package “fgsea”^61^, with gene sets obtained from MSigDB^47^, using parameters “minSize=0”, “maxSize=10000” and “nperm=10^7^”. Categories with *q* < 0.05 were considered statistically significant.

### RT-qPCR validation of genes identified from differential expression analysis

RNA was extracted from the indicated cell lines with Nucleospin RNA II kit (Macherey-Nagel), and converted to cDNA using iScript cDNA synthesis kit (Bio-Rad). qPCR was performed using 2x qPCR SyGreen Mix (PCR Biosystems) and the CFX Connect Real-Time System (Bio-Rad). Data analysis was performed by comparing ΔΔ CT values using Ubiquitin as a reference gene.

### Transcriptome comparison with tumors from TCGA

RNA sequencing data for 9,583 tumors from 32 cancer types were downloaded from the cgHub repository on December 18, 2015, and aligned to the hg19 human genome assembly, excluding alternative haplotype regions, with hisat^62^ 0.1.6-beta (parameters: “--no-mixed -- no-discordant --no-unal --known-splicesite-infile”), using splice junctions defined in the GENCODE (version 19) reference human genome annotation. Gene read counts were derived with htseq-count^54^ (parameters: “-m intersection-strict -s no”). RPKM normalized values were calculated, taking into account the max mature transcript length of each gene and using robust size factors as previously described for the DESeq method^63^. For the correlation analysis, reads from our own sample was realigned and read counts requantified and normalized using the same methods described for TCGA data. However, standard read depth-based size factors were used for the RPKM normalization of this sample. Pairwise Spearman correlation coefficients were then calculated between our sample and each TCGA sample, with respect to all coding genes (using the function “corr” in MATLAB R2018a). For t-distributed stochastic neighbor embedding (t-SNE) analysis, log_2_ transformed (pseudocount of 1 added) expression values of all coding genes were used, together with the “Rtsne” function from the “Rtsne” R package^64^. A separate classification was performed using a 6-nearest neighbor approach based on Spearman correlations, as previously described^65^.

### Copy number segmentation and purity estimation

Copy number segmentation was performed using binocular (https://sourceforge.net/projects/binocular), with input from an unfiltered VCF file from MuTect for a given sample, together with WGS BAM files for tumor and normal samples. Parameters used were “--delta=90,” “--min-maf-delta=0.05,” “--ai-cutoff=0.001” and “--min-copy-ratio=1.1” for the majority of samples, although for samples with more variable coverage this threshold was raised. For tumors without matching normals, the intersect of segments defined using normals from the other samples were used. Sample purity and ploidy was estimated with ichorCNA^66^, using the parameters “--ploidy “c(2,3,4)” --normal “c(0.1,0.2,0.3,0.4,0.5,0.6,0.7,0.8,0.9)” --maxCN 10“, based on the segmented copy number values.

### Associations between broad copy number changes and metastatic or primary tumors

Segmented copy number data from TCGA primary tumors were downloaded from GDC Data Portal (accessed on 6 October 2017). Copy number changes with an absolute log_2_ ratio relative to diploid chromosomes less than 0.2 and with width less than 10^6^ base pairs were filtered out from both TCGA UMs and our tumors. The general events to test were defined as those where a contiguous altered region spanning all events in all metastasis samples were present that had a width of at least 10^6^ base pairs, and which occurred in at least 5% of samples in either dataset, to avoid events that were unlikely to be relevant to selection. Changes of same direction (loss or gain) affecting each region were then assessed for association with each of the two datasets using Fisher’s exact test (two-tailed) using the function “fisher.test” in R 3.5.0. Since the resulting contiguous regions practically spanned the entire length of each affected chromosome arm, with the exception of chromosome 3, which spanned the entire chromosome, one test was performed per gain or loss event of each such arm. *p*-values were corrected for multiple testing using the Benjamini-Hochberg method.

### Ranking of genes in broad copy number aberrations

RNA-seq data for the TCGA UVM dataset (*n* = 80) were downloaded using the TCGAbiolinks R package^67^, with parameters “project = ‘TCGA-UVM’, data.category = ‘Transcriptome Profiling’, data.type = ‘Gene Expression Quantification’, workflow.type = ‘HTSeq - Counts’”. Read counts were normalized using the “rpkm” method from the “edgeR” package (“log=FALSE, prior.count=1”), with gene lengths chosen as the maximum transcript length obtained via biomaRt and the “ensembl” database. Segmented copy number profiles for each sample were downloaded from the GDC data portal. The copy number status of each gene was calculated by choosing the maximal absolute log_2_ ratio among segments spanning the gene. Genes with both copy number and gene expression values assigned were retained.

To focus on genomic regions subject to copy number changes recurrent enough to indicate selection, TCGA GISTIC results^2^ were used (obtained from gdac.broadinstitute.org, accessed 4 July 2017). Recurrent broad copy number changes with *q*-values < 0.05 were retained. To focus on genes that were altered at relevant frequencies and more likely to be part any minimal region of overlap, genes with an absolute log_2_ copy number ratio < 0.2 were filtered out and only genes with an alteration frequency in the upper third quartile per chromosome arm event were retained. The third quartile was chosen, rather than a stricter threshold, since some regions may be subject to low-frequency focal events of a random or artifactual nature.

To find genes altered in expression in tandem with the copy number changes, linear regression between copy number and expression was performed, adjusting for tumor purity estimates obtained from Zheng et al.^68^. Genes with too low expression variance to test were removed (defined as those for which regression failed to converge). Univariate survival tests with Cox regression (the “coxph” function from the “survival” R package^69^) were then carried out against clinical data downloaded using TCGAbiolinks^67^, using the variables “vital_status”, “days_to_death” and “days_to_last_follow_up”.

Segmented copy number values from our metastasis samples were mapped to gene names as described above and converted to log_2_ ratios. Values of zero prior to transformation were set to the lowest observed non-zero copy number value. Gene expression values for the metastasis dataset were normalized with RPKM as described above and batch corrected using the “removeBatchEffect” function of the “limma” R package^70^. Genes with values in both the copy number and gene expression dataset as well as preserved after the filtering on the TCGA data were retained. Associations between gene expression and copy number status were assessed as for the TCGA dataset, considering sample purity. *p*-values from associations in the TCGA data and metastasis dataset were combined using Fisher’s method and FDR adjusted using the Benjamini-Hochberg method. Candidates with *q*-values < 0.05, an independent raw *p*-value of less than 0.05 in each dataset and correlations consistent with the direction of the assessed copy number event were retained. Candidates in regions with more samples harboring gains than losses were retained as candidates of gains and vice versa.

To assess the extent to which a given gene may have a wider impact on cellular behavior, manually curated protein-protein interactions with experimental evidence defined in the Human Protein Reference Database (HPRD)^71^ were used. The database was accessed using the “iRefR” R package^72^ and node degrees calculated using the “degree” function in the “igraph” package^73^. The candidates were then ranked by the number of HPRD connections, and then by whether any univariate survival associations existed (p < 0.05) implying worse survival consistent with the nature (gain or loss) of the copy number event assessed. This way, survival associations were placed a low weight, with the motivation that such associations are easily confounded by multiple genomic and clinical factors.

### siRNA screen

*In vitro* knockdown of selected genes was performed using siRNA in UM22 cells. Transient transfection was performed with mock siRNA (control), a positive control siRNA (GNAQ) or a pool of 4 siRNA per gene of interest. The siRNA duplexes were purchased from Dharmacon (Thermo Fisher Scientific, Waltham, MA, USA) and the lipid based transfection was performed with Lipofectamine-RNAiMAX® (Thermo Fisher Scientific, Waltham, MA, USA) using 1 pmol of siRNA per well of a 96-well plate as per the guide line provided by manufacturer. The RNA-Lipid complex was made in Opti-MEM® Reduced Serum medium. The cells were seeded in black 96-well plates (Corning) and 72 hours post-transfection cells and viability of was monitored with ATP measurement using CellTiter-Glo® Assay (Promega) the luminescence was measured with GloMax Discover plate reader (Promega). In parallel the manual cell count was performed using Trypan blue staining of cells obtained from transfections in 12-well format.

### Single-cell RNA-seq analysis of immune infiltrates

Small pieces of tumor biopsies were cultured for two weeks in RPMI medium containing 10% human serum and 6000 U/ml IL-2. Young TIL (yTIL) cultures were then cryopreserved before use. Two days before performing the single cell experiments, yTIL cultures were thawed. Cells were counted and 7 000 cells were injected into a single cell library preparation instrument (10x Genomics). The steps following were performed using the Single Cell V(D)J kit according to the kit description (10x Genomics). V(D)J libraries were sequenced on a MiSeq instrument (Illumina) whereas the gene expression libraries were run on a NextSeq (Illumina). Single-cell transcriptomics data were aligned against the hg38 reference genome and preprocessed using the Cellranger pipeline provided by 10x Genomics. Expression levels were estimated using the Cellranger “count” function, with default parameters. TCR chain assembly was also performed using the Cellranger pipeline, using default parameters. t-SNE maps for each sample were generated using the SingleR R package (version 1.0) and cell types were inferred using the approach described by Zheng et al.^49^, with two modifications: correction of a code error that misclassified some CD4+ cells and reclassification of cells classified as memory CD4+ cells not expressing CD4 but CD8 as the closest matching non-CD4+ cell type. Doublet cells were defined as those expressing more than one alpha or beta chain and those that were classified as doublets by the tool DoubletFinder^74^ (default parameters). TCR clonotype diversity was assessed for CD8+ cells using clonotype frequency and the “diversity” function (type = “e”) from the “diverse” R package^75^. Clonotype activation states were defined by first independently clustering each clonotype by Spearman correlations with respect to the mean expression of cytotoxicity markers and T cell terminal differentiation / exhaustion markers obtained from Azizi et al.^76^, using complete linkage hierarchical clustering with a Euclidean distance metric (“pheatmap” R package). The number of clusters were defined based on the number of clusters maximizing the clustering indices calculated with the NbClust R function (“NbClust” package; parameters: min.nc=4, max.nc=20, method = “complete”, index = index[i], alphaBeale = 0.1, where index[i] indicates each of the supported indices for a distance matrix). The resulting clusters were then used to define overlapping sets of clusters.

### Flow cytometry

Single cell suspensions from cryopreserved tumor biopsies and yTILs were surface stained for 30 minutes in RT. The following antibodies were used for surface staining: CD3 (HIT3a), CD4 (A161A1), CD8 (HIT8a), CD45 (2D1), CD69 (FN50), CTLA-4 (BNI13), PD-1 (EH12.H7), TIGIT (A15153G) and TIM-3 (F38-2E2) from BioLegend and CD39 (eBioA1) from eBioscience. For detection of melanoma antigen specific CD8 T cells, cells were surface stained for 45 min in 37°C using Melanoma Dextramer Collection 1 kit from Immudex. Dead cells were excluded from the analysis using Live/Dead Aqua (Invitrogen). Flow cytometry data was acquired using BD Accuri C6 (BD Biosciences), BD LSRFortessa X-20 (BD Biosciences) or BD FACSARIA FUSION (BD Biosiences) and analysed using FlowJo software (FlowJo LLC). The gating strategy is shown in **Fig. S6**.

### Ethics

The patients received oral and written information and signed the informed consent agreement according to the ethical approval at the Regional ethical review board (#289-12 and # 144-13). Animal handling: Regional animal ethics committee of Gothenburg approval #36-2014.

## Acknowledgements

The results published here are in part based upon data generated by the TCGA Research Network (cancergenome.nih.gov/). We thank Sofia Stenqvist for animal care, Ola Nilsson and Gülay Altiparmak for histology, Carina Karlsson for technical assistance and Therese Bengtsson and Valerio Belgrano for patient registrations. The GeneCoreSU and SciLifeLab facilities are acknowledged for sequencing services, partly financed by a National Genomics Initiative grant to JAN. Other funding sources included Knut and Alice Wallenberg Foundation, Cancerfonden, Vetenskapsrådet, Familjen Erling Perssons stiftelse, and Västra Götalandsregionen.

## Supplementary Figure Legends

**Figure S1:** Multiple *BAP1* mutations in UM16 present in both metastasis and primary tumor.

**Figure S2:** Tumor purity and ploidy analysis. **a)** Tumor purity per sample. **b)** Tumor ploidy. Values were estimated using ichorCNA.

**Figure S3: a)** *GNA11* Q209L mutation and **b)** *BAP1* deletion in UM11 DNA and RNA. **c)** Transcriptomic classification of UM11 using t-SNE against TCGA tumors (n = 9,583) from 32 cancer types. 6-nearest neighbor classification based on Spearman correlation coefficients, according to a previously described approach45, gave that 6/6 of the top ranked samples in TCGA were UMs (average correlation coefficient 0.93). **d-e)** Clinical manifestation of an iris melanoma. At diagnosis an iris nevus was seen (c) which progressed to an iris melanoma (d) two years later. e-g) Histological sections of different magnifications showing the locally invasive iris melanoma.

**Figure S4:** Focal deletions of *CDKN2A***. a)** RNA-seq reads of the *CDKN2A* locus in UM9 and UM22 metastases and PDX models. **b)** Exome sequencing of tumor-infiltrating lymphocytes confirming the somatic identity of *CDKN2A* deletion in UM22.

**Figure S5:** Immunohistochemistry of PDX models with respect to hematoxylin and eosin, Melan-A, HMB-45 and S100-P.

**Figure S6:** Estimated number of neoepitope-generating mutations per sample, using netMHCpan, with HLA genotype inferred using polysolver.

**Figure S7:** Gating strategy for identification of CD4+ and CD8+ T cells among REP-TILs, yTILs and original material from the metastasis. **a)** Representative plots from UM13 showing the gating strategy used to identify MART-1 specific CD8+ T cells among REP-TILs. **b)** Gating strategy from yTIL material of UM22 illustrating the gating strategy for analysis of CD4+ and CD8+ T cells in PR and yTIL samples. **c)** Flow cytometry analysis of T-cells, with respect to CD4 and CD3. **d)** Single-cell RNA expression levels of *CD3G*, *CD8A*, *CD4* and *NCAM1* in TILs.

**Figure S8:** Analysis of T cell receptor clonotypes. **a)** Clustering of CD8+ T-cell clonotypes using average clonotype expression and Spearman correlation, based on exhaustion / terminal differentiation genes and cytotoxicity genes. **b)** Clonotypes in the intersection from the clusters in (a), colored by membership in the clusters from (a). Sizes of dots are proportional to the number of cells having each clonotype.

**Figure S9:** Flow cytometry analysis of T-cells. Proportions of **a)** CD8+ and **b)** CD4+ cells positive for PD-1 and TIGIT. Proportions **c)** CD8+ and **d)** CD4+ cells positive for CTLA-4.

**Figure S10:** Flow cytometry analysis of T-cells. Proportions of **a)** CD8+ cells and **b)** CD4+ cells positive for PD-1 and TIM-3.

## Supplementary Table Legends

**Table S1:** Clinical details of patients.

**Table S2:** Annotated mutations detected in each tumor.

**Table S3:** Statistics for tests of association between broad copy number changes in our samples versus TCGA tumors.

**Table S4:** Ranking of genes relevant in broad copy number changes, per affected chromosome arm. Genes are ordered by presence of correlation between copy number and expression in both datasets, known protein-protein interactions and univariate survival statistics.

**Table S5:** Enriched Reactome gene sets among genes in the combined ranked lists either regions of gain or loss.

**Table S6:** Differentially expressed genes between biological replicates (*n* = 3 per condition) transduced with functional *BAP1* vectors or empty vectors.

**Table S7:** Gene set enrichment analysis for the category “chemical and genetic perturbations” of MSigDB of genes assessed in the comparison of replicates transduced with functional *BAP1* vectors or empty vectors.

