## Supplementary figures and images for "Molecular profiling of driver events and tumor-infiltrating lymphocytes in metastatic uveal melanoma"

### Figure S1

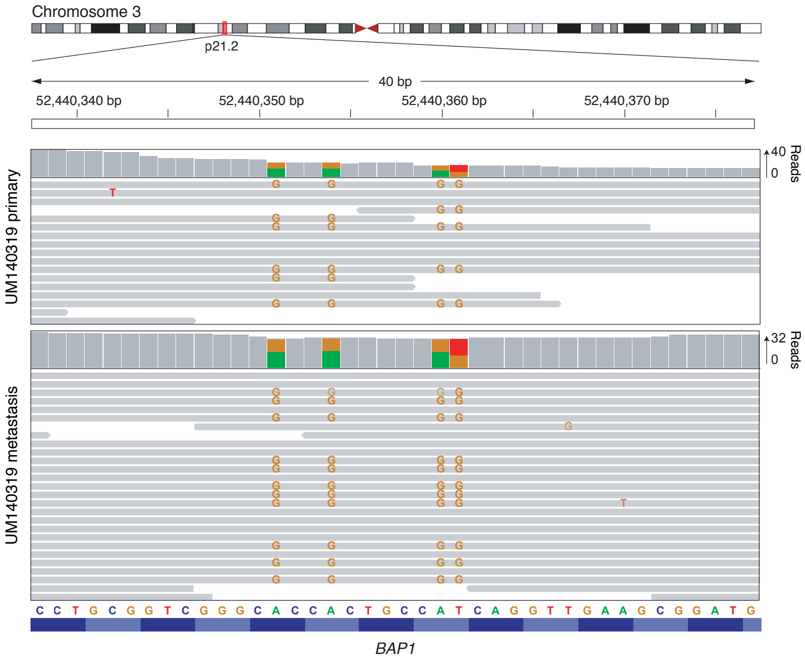

### Figure S2

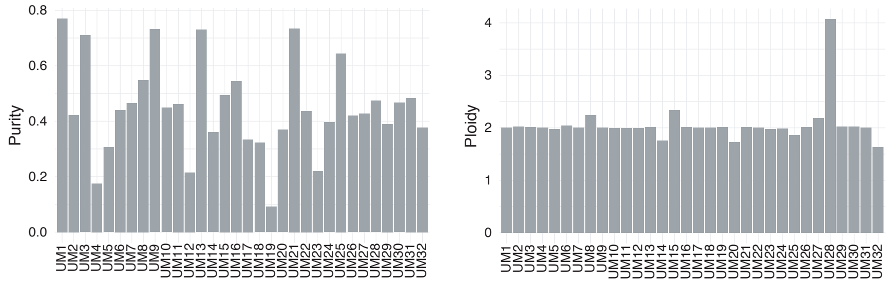

### Figure S3

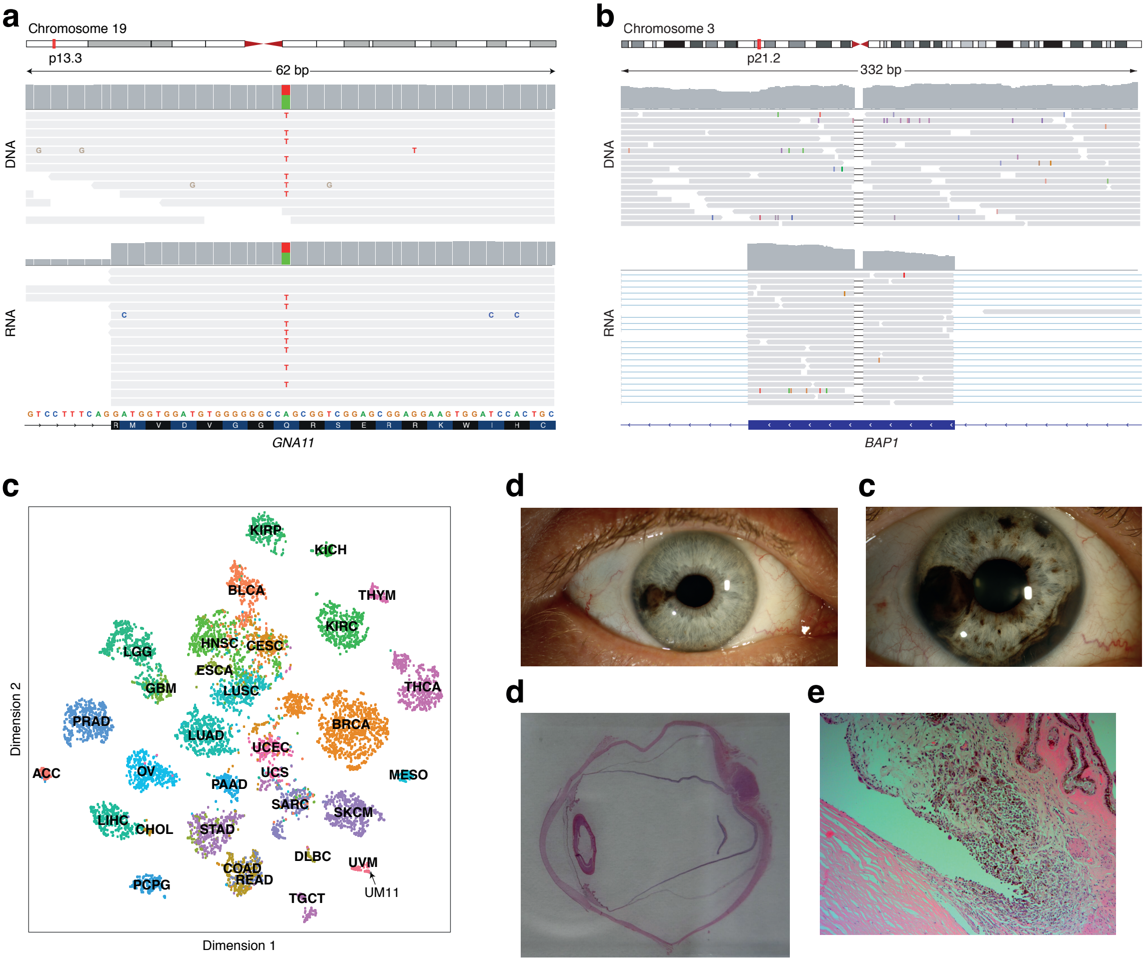

### Figure S4

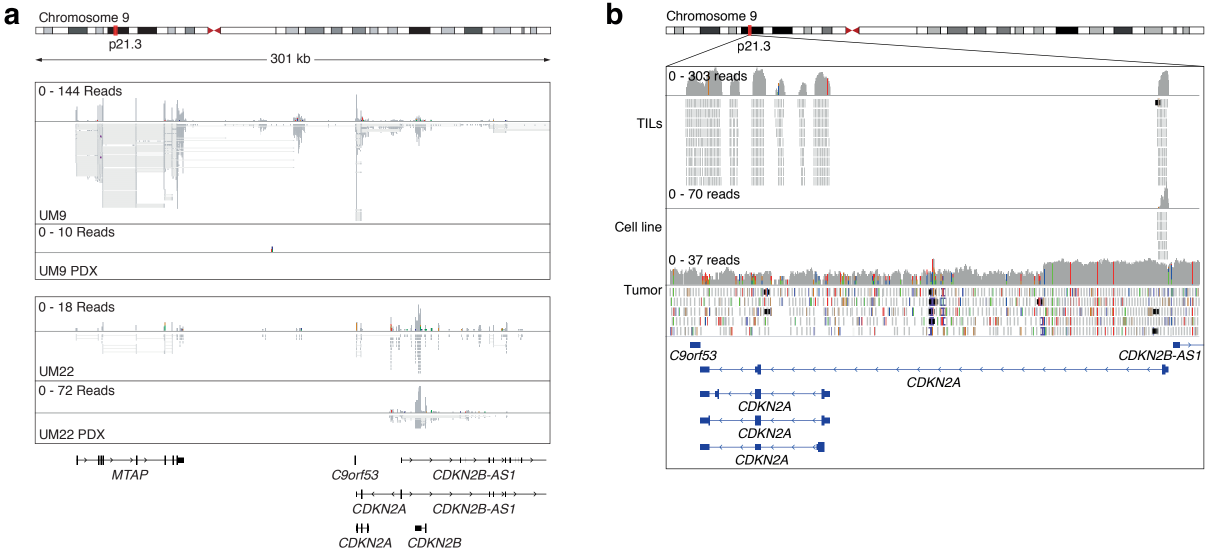

### FIgure S5

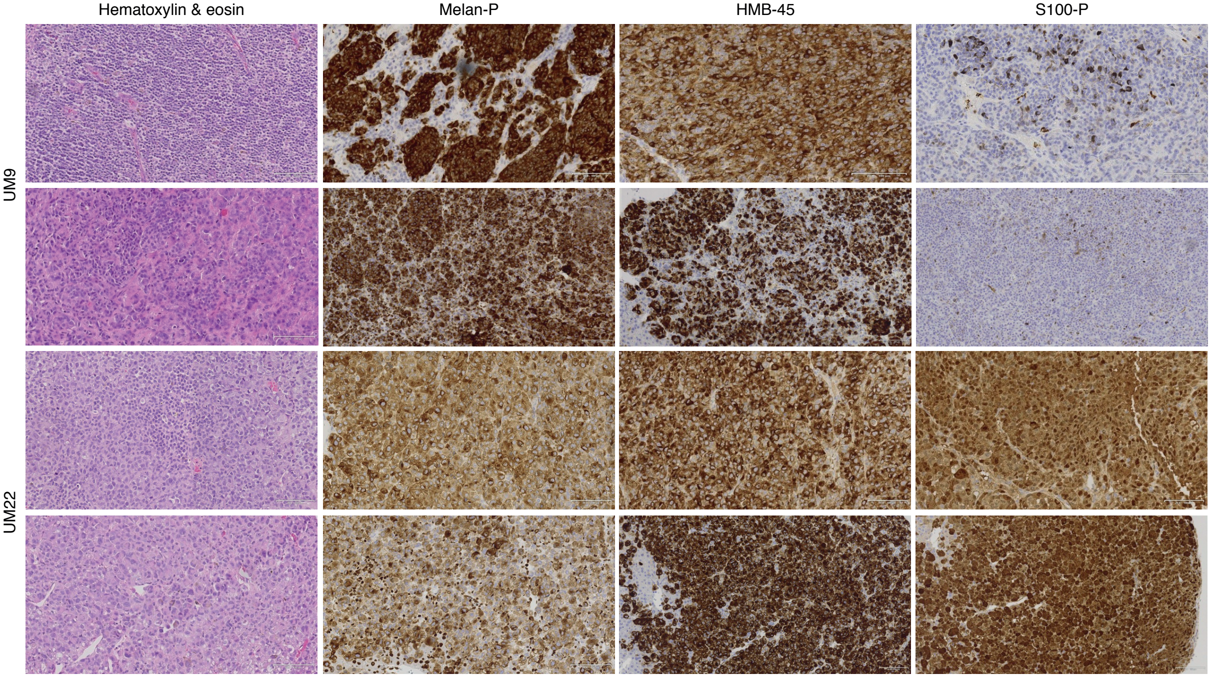

### Figure S6

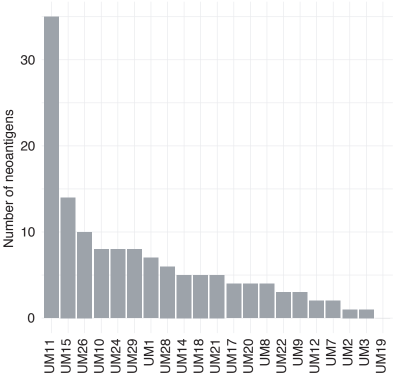

### Figure S7

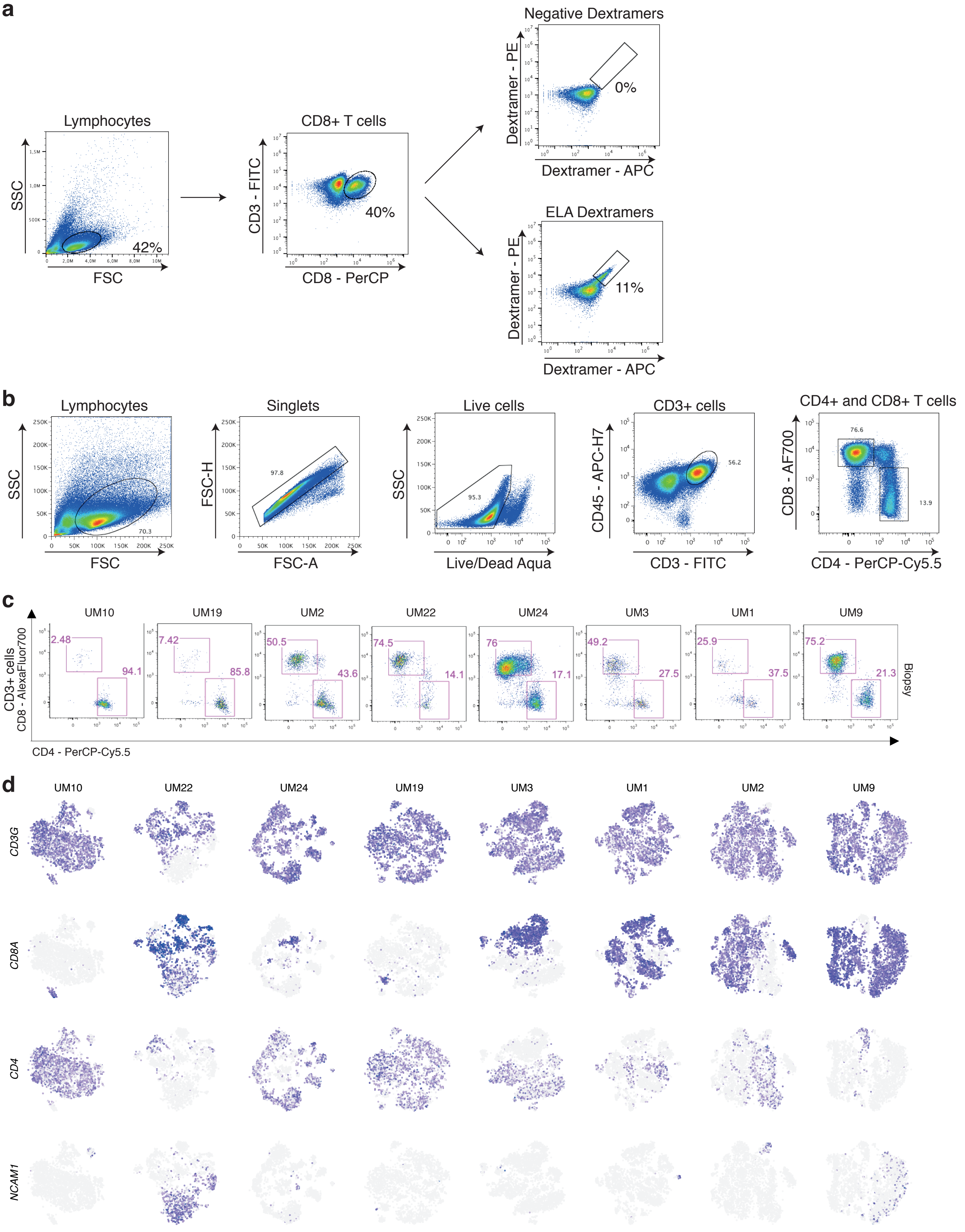

### Figure S8

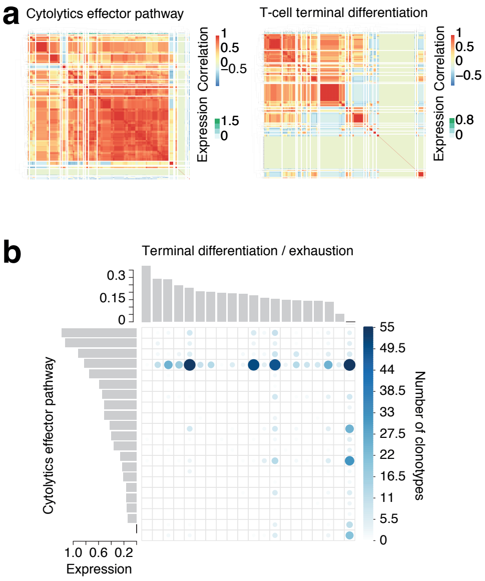

### FIgure S9

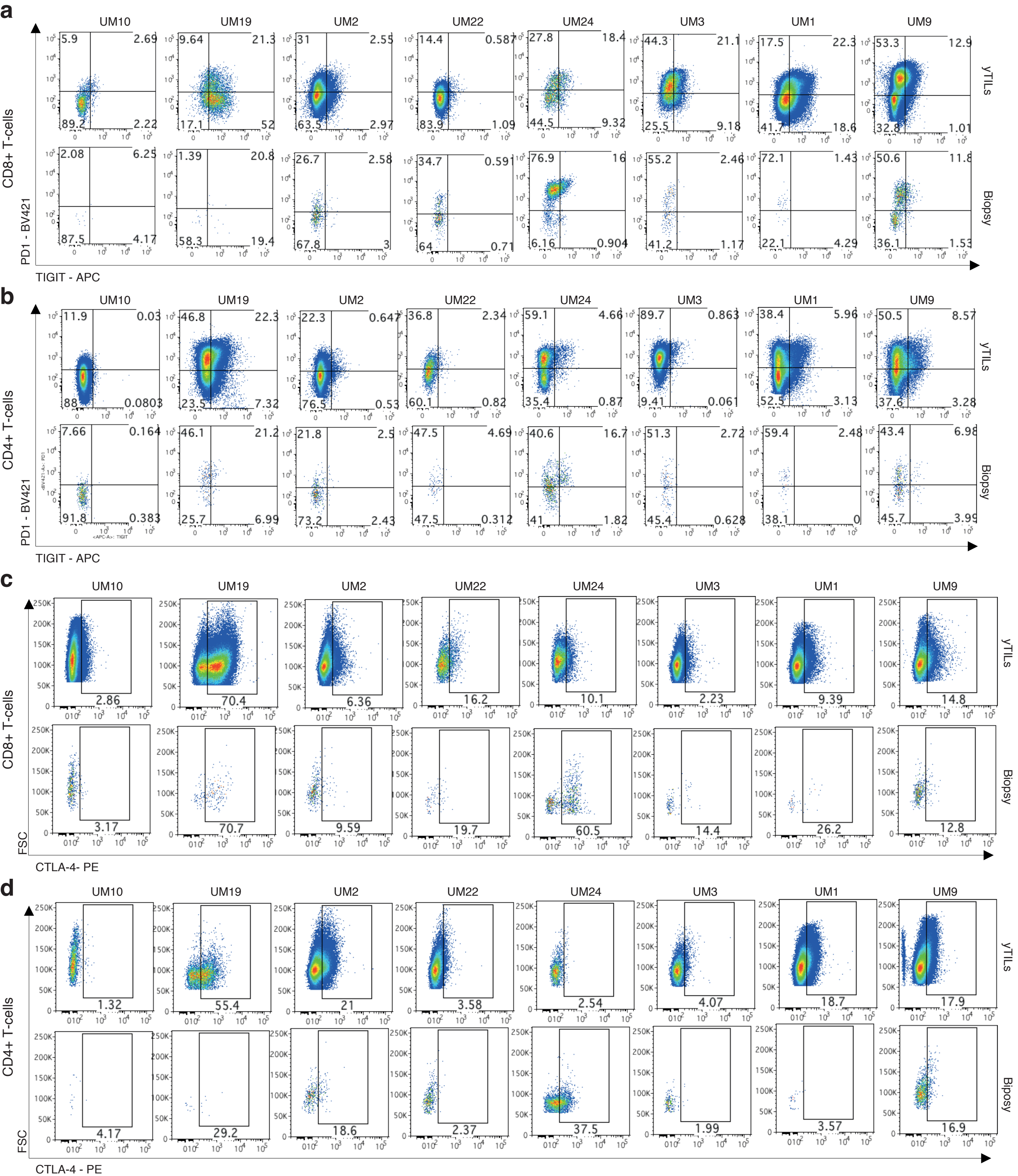

### FIgure S10

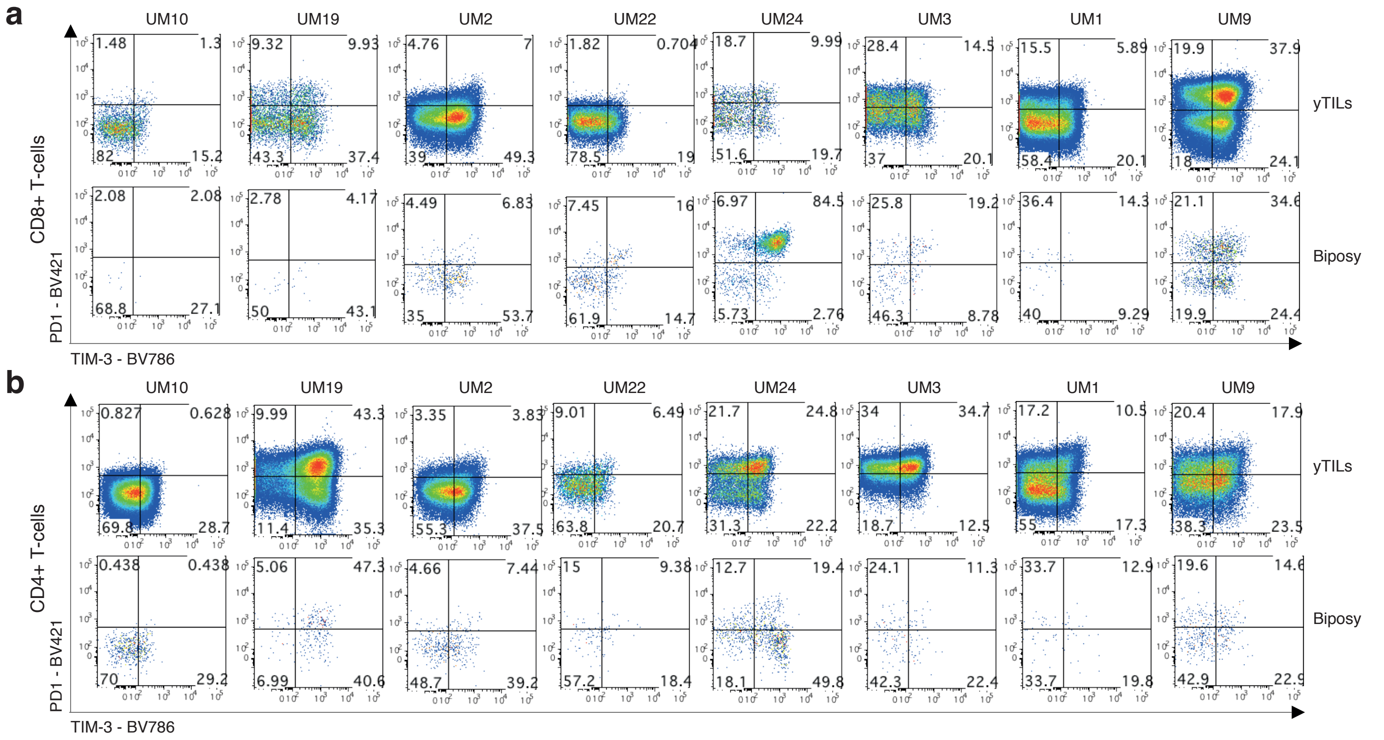
